# Phylogeny Reveals Novel HipA-Homologous Kinase Families and Toxin – Antitoxin Gene Organizations

**DOI:** 10.1101/2021.04.14.439929

**Authors:** Kenn Gerdes, Rene Bærentsen, Ditlev E. Brodersen

## Abstract

Toxin – Antitoxin modules function in the genetic stability of mobile genetic elements, bacteriophage defense, and antibiotic tolerance. A gain-of-function mutation of the *Escherichia coli* K-12 *hipBA* module can induce antibiotic tolerance in a subpopulation of bacterial cells, a phenomenon known as persistence. HipA is a Ser/Thr kinase that phosphorylates and inactivates glutamyl tRNA synthetase, inhibiting cellular translation and inducing the stringent response. Additional characterized HipA homologues include HipT from pathogenic *E. coli* O127 and YjjJ of *E. coli* K-12, which are encoded by tri-cistronic *hipBST* and monocistronic operons, respectively. The apparent diversity of HipA homologues in bacterial genomes inspired us to investigate overall phylogeny. Here we present a comprehensive phylogenetic analysis of the Hip kinases in bacteria and archaea that expands on this diversity by revealing seven novel kinase families. Kinases of one family, encoded by monocistronic operons, consist of an N-terminal core kinase domain, a HipS-like domain and a HIRAN (HIP116 Rad5p N-terminal) domain. HIRAN domains bind single or double-stranded DNA ends. Moreover, five types of bicistronic kinase operons encode putative antitoxins with HipS-HIRAN, HipS, γδ-resolvase or Stl repressor-like domains. Finally, our analysis indicates that reversion of *hipBA* gene-order happened independently several times during evolution.

**Importance:** Bacterial multidrug tolerance and persistence are problems of increasing scientific and medical significance. The first gene discovered to confer persistence was *hipA*, encoding the kinase toxin of the *hipBA* toxin-antitoxin (TA) module of *E. coli*. HipA-homologous kinases phosphorylate and thereby inactivate specific tRNA synthetases, thus inhibiting protein translation and cell proliferation. Here, we present a comprehensive phylogenetic analysis of bacterial Hip kinases and discover seven new families with novel operon structures and domains. Overall, Hip kinases are encoded by TA modules with at least 10 different genetic organizations, seven of which have not been described before. These results open up exciting avenues for the experimental analysis of the superfamily of Hip kinases.

## Introduction

Prokaryotic toxin-antitoxin (TA) modules were discovered due to their ability to stabilize plasmids by killing of plasmid-free cells by a mechanism known as *post-segregational killing* (1, 2). The mechanism relies on stable protein toxins that are inhibited either by unstable antitoxin RNAs (type I and III TAs) or unstable antitoxin proteins (type II TAs) as long as the plasmid remains in the cell. If, on the other hand, the plasmid is lost, degradation of antitoxin leads to toxin activation and hence, death of plasmid-free cell. Since their discovery on plasmids, TAs have been identified on a wide range of bacterial and archaeal chromosomes as well (3–5), often in multiple or even large numbers (5–9). For example, *Mycobacterium tuberculosis* encodes ~70 type II TA modules (7, 10, 11) while the plant symbiont *Sinorhizobium meliloti* contains more than 100 (12). The biological functions of chromosome-encoded TAs are debated but experimental evidence supports at least three roles, that are not mutually exclusive: (i) genetic stabilization of chromosome segments or entire chromosomes (13–16); (ii) anti-phage defense by abortive-infection (17, 18), and (iii) antibiotic tolerance (19–22). Intriguingly, it was recently discovered that bacterial retrons encode a type of three-component TAs that can function as anti-phage defense systems thus supporting the notion that a major function of TAs may in fact be to curb or control bacteriophage infection (23–26). This idea is consistent with the recent observation that environmental or nutritional stress in general does not activate Type II TA-encoded toxins (27).

Evidence supports that some TAs induce antibiotic tolerance or persistence in bacteria. Persistence is a phenomenon found in all bacteria tested (19, 28–30) and is operationally defined the subpopulation of a bacterial cells that survive for an extended period of time in the presence of inhibitory concentrations of antibiotics (31). Common to persistence mechanisms is that the phenotype is a stochastic phenomenon and only expressed by a fraction of the cell population at any given time (31, 32). Importantly, persistence is believed to contribute to the recalcitrance of bacterial infections and may thus pose a significant medical problem (30, 33–35). At the mechanistic level, persisters are slow-growing cells that display increased survival rates in the presence of antibiotics (32, 36, 37). In addition, this also buys the bacterial population time to develop true antibiotic resistance (38). Apart from a reduced growth-rate, persister cells can also arise from expression of high levels of factors that counter the effects of antibiotics in a small subpopulation of cells (39, 40).

*hipA* (<u>h</u>igh <u>p</u>ersister <u>g</u>ene <u>A</u>) of *Escherichia coli* K-12 was the first gene found to be associated with increased persistence based on the identification of the gain-of-function allele *hipA7* in a strain exhibiting increased tolerance towards penicillin (41). The mutant allele, later also found in clinical isolates of uropathogenic *E. coli* (28), exhibits a 100 to 1,000-fold increased level of persistence (32, 42), but even the *wt hipBA* module can be shown to confer a modest, but measurable, increase of persistence (28). The *hipA* toxin gene and its upstream *hipB* antitoxin gene constitute a canonical type II TA module encoding two proteins that combine to form an inactive HipBA complex, which, upon degradation of HipB, generates active HipA toxin (**Figure 1A**) (43). Consequently, ectopic production of HipA in *E. coli* causes severe growth inhibition that can be reversed by later expression of HipB antitoxin (43). HipBA from both *Escherichia coli* K-12 and *Shewanella oneidensis* MR-1 assemble into hetero-tetrameric <u>HipA_2_B_2_ complexes</u> (28, 44–48).

**Figure 1.**
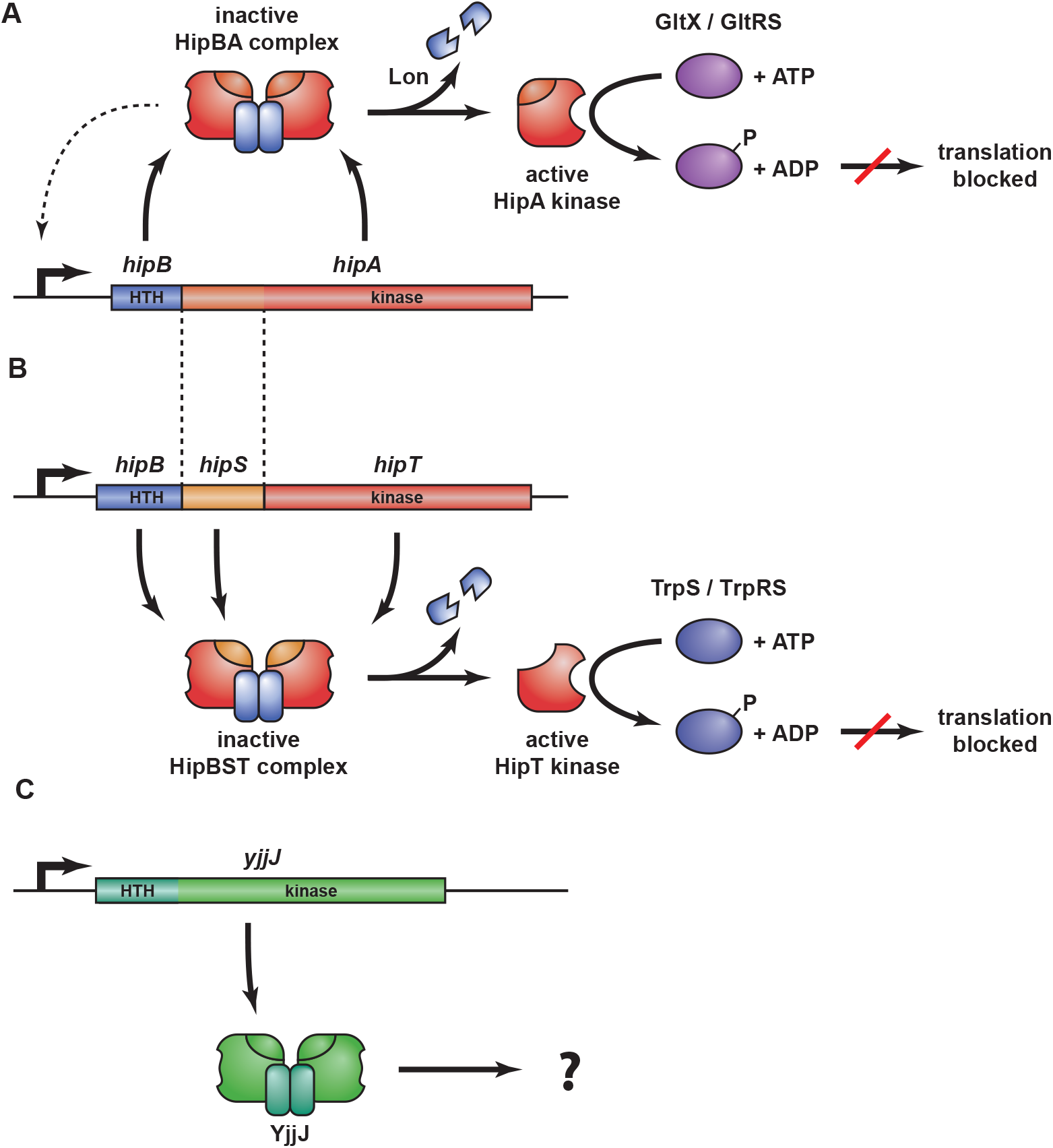
Overview of the *hipBA*, *hipBST*, and *yjjJ* operons and their protein products. (**A**) *hipBA* encodes the antitoxin HipB (blue) and toxin kinase HipA (red/orange) that form an inactive HipB_2_HipA_2_ complex that can bind to operators in the promoter region via HTH DNA binding domains in HipB and thereby autoregulate transcription (dashed line) (28). Upon HipB degradation by Lon protease (88) and activation of HipA, HipA phosphorylates and inhibits glutamine tRNA synthetase (GltX/GltRS), thereby halting translation and inducing the stringent response (51, 52, 56). (**B**) *hipBST* encodes three proteins, HipB (blue), HipS (orange), and HipT (red) that form an inactive HipBST complex (59). HipS is homologous to the N-subdomain-1 of HipA and functions as the antitoxin that neutralizes HipT (59). Like HipB of HipBA, HipB of HipBST has a HTH domain and augments the inhibition of HipT by HipS but does not function as an antitoxin on its own. Free HipT phosphorylates and inhibits tryptophan tRNA synthetase (TrpS) and thereby halts translation in a similar fashion as HipA. (**C**) *yjjJ* is a single cistron operon that encodes a HipA-homologous kinase YjjJ (green) that, when overproduced, inhibits cell growth (60). YjjJ, that we coin HipH, has an HTH-domain in its N-terminus that may function to bind DNA.

HipA is a so-called Hanks serine-threonine kinase (49, 50) that specifically targets a conserved serine residue (Ser^239^) in glutamyl-tRNA synthetase (GltX) inside its bacterial host, inhibiting the enzyme and thereby aminoacylation of tRNA^Glu^ (51, 52). As a consequence, the ratio of charged to uncharged tRNA^Glu^ decreases, thus stimulating RelA-tRNA^Glu^ complexes to bind the ribosomal A site. Activation of RelA (53) on the ribosome leads to an increase in the cellular (p)ppGpp level triggering the stringent response (42, 51, 54). HipA shares its fold with human Cyclin Dependent Kinases and maintains all of the conserved motifs necessary for kinase activity (45). The antitoxin HipB contains a classical (Cro-like) helix-turn-helix (HTH) DNA-binding domain (45) and forms a homodimer when in complex with HipA that allows binding to palindromic operator sequences in the *hipBA* promoter region (**Figure 1A**). The mechanism of toxin inhibition by HipB has not been fully elucidated, but it appears to differ from most type II systems in that the antitoxin does not interact directly with the toxin active site. Instead, the very C terminus of HipB appears to bind in a pocket on HipA, possibly regulating toxin activity (44). Finally, HipA can inactivate its own kinase activity by auto-phosphorylation (55), a phenomenon that has been proposed to function in resuscitation of persister cells (46, 56).

HipA homologues are present in many bacteria. For example, the chromosome of the alfaproteobacterium *Caulobacter crescentus* contains three *hipBA* loci for which the encoded HipA toxins inhibit protein synthesis upon ectopic expression in all three cases (57). Like their *E. coli* counterpart, HipA_1_ and HipA_2_ contribute to persistence during stationary phase by phosphorylating aminoacyl-tRNA synthetases *in vivo*: HipA_2_ targets lysyl-tRNA synthetase (LysS), tryptophan-tRNA synthetase (TrpS), and GltX, while HipA_1_ phosphorylates TrpS and GltX (57). In both cases, the stringent response regulator SpoT (Rel) is required for *hipBA*_1_ and *hipBA*_2_-mediated persistence. A recent report confirmed that HipA_2_ only phosphorylates and inhibits TrpS whereas LysS and GltX were not found to be phosphorylated (58). Nevertheless, both studies agreed that HipA_2_ induces the stringent response and persistence in parallel (57, 58).

We recently described a new family of minimal bacterial kinases, HipT, members of which exhibit sequence similarity with the C-terminal part of HipA but are encoded by three-gene operons (**Figure 1B**) (59). HipT of *E. coli* O127 is functionally similar to HipA and targets tryptophanyl-tRNA synthetase (TrpS/TrpRS) at two conserved serine residues, inactivating the enzyme. Likewise, ectopic production of *hipT* inhibits cell growth and translation and, consistently, stimulates production of (p)ppGpp (59). The gene immediately upstream of *hipT* encodes a small protein, HipS, that exhibits sequence similarity to the N-terminal part of the larger HipA kinase (**Figure 1B**). Surprisingly, HipS neutralizes HipT *in vivo*, and therefore appears to function as the antitoxin of the *hipBST* module. The third component, HipB, encoded by the first gene of the *hipBST* operon (**Figure 1B**), contains an HTH domain and is homologous to HipB of HipBA but does not counteract HipT kinase activity directly. Rather, this protein functionally appears to augment the ability of HipS to neutralize HipT (59). The structural and mechanistic details that set the bi-cistronic and tri-cistronic Hip kinase systems apart have not yet been elucidated. Finally, *E*. *coli* K-12 encodes the HipA homologue YjjJ in a monocistronic operon, thus lacking an adjacent antitoxin gene, which also inhibits cell growth upon induction (**Figure 1C**) (60). Interestingly, YjjJ contains a HTH domain at its N terminus that may compensate for the lack of a DNA-binding antitoxin; however, how YjjJ kinase activity is controlled, remains unknown.

The discovery of the *hipBST* tri-cistronic operons (59) as well as the observed diversity among Hip toxin homologues in various bacterial species inspired us to investigate the overall phylogeny of HipA-homologous proteins and their gene families in prokaryotic microorganisms. This led to the discovery of seven novel Hip kinase families, potential antitoxins with novel features such as HIRAN (HIP116 and RAD5 N-terminal) domains with predicted specificity for single-stranded or double-stranded DNA ends and a novel putative two-domain antitoxin family consisting of a HipS-like domain coupled to a HIRAN domain. We also find evidence that HipA-homologous kinases are present in Archaea. Together, these results delineate the structural and functional diversity of the family of HipA kinases and suggest directions for future experimental research.

## Results and Discussion

### HipA-homologous kinases form a strongly supported, bifurcated phylogenetic tree

The phylogenetic analysis was initiated using HipA and YjjJ of *E. coli* K-12 and HipT of *E. coli* O127 as seed sequences using BLASTP and HMMSEARCH (see *Methods* for details). While this revealed a vast number of HipA-homologous kinases within the bacterial domain and a few in the archaeal domain, searches in the Eukarya domains did not disclose significant homologues (E > 10^−5^). To analyze the vast number of high-score homologues (E > 10^−10^) systematically, we repeated the search using individual bacterial and archaeal phyla as search spaces (**Figure S1**) and retrieved ~1,800 high-scoring Hip homologues. Tenacious curation, including manual inspection of putative neighboring antitoxin genes of each individual kinase gene, removal of incomplete genes and exclusion of closely related kinases (< 5% sequence difference), reduced the number of included kinases to a final set of 1,239 sequences. The majority of these are from the phylum Proteobacteria (70%) while they are also frequently observed among Actinomycetes (13%), Firmicutes (5%), and Spirochetes (2%) (**Table S1**). Using these sequences, we generated a phylogenetic tree, called the “Hip Tree” (**Figure 2A**). Fully annotated and bootstrapped versions of the Hip Tree are shown in **Figures S2A and S2B** while a full breakdown of the phylogeny of the kinases is given in **Table S2**A. The Hip Tree consists of 11 Main Clades (Clades I-XI) all supported by high statistical confidence levels (bootstrap values ≥ 98%). The tree is bifurcated with one branch containing Main Clades I through X while the other branch consists of the diverse Main Clade XI. In **Figure 2A**, the colours of the clades (shown as triangles) reflect kinases encoded by TA modules with identical genetic organization, as explained in the next section. In other words, each differently colored triangle in **Figure 2A** reflects HipA-homologous kinases encoded by TA modules with a different genetic organization.

**Figure 2.**
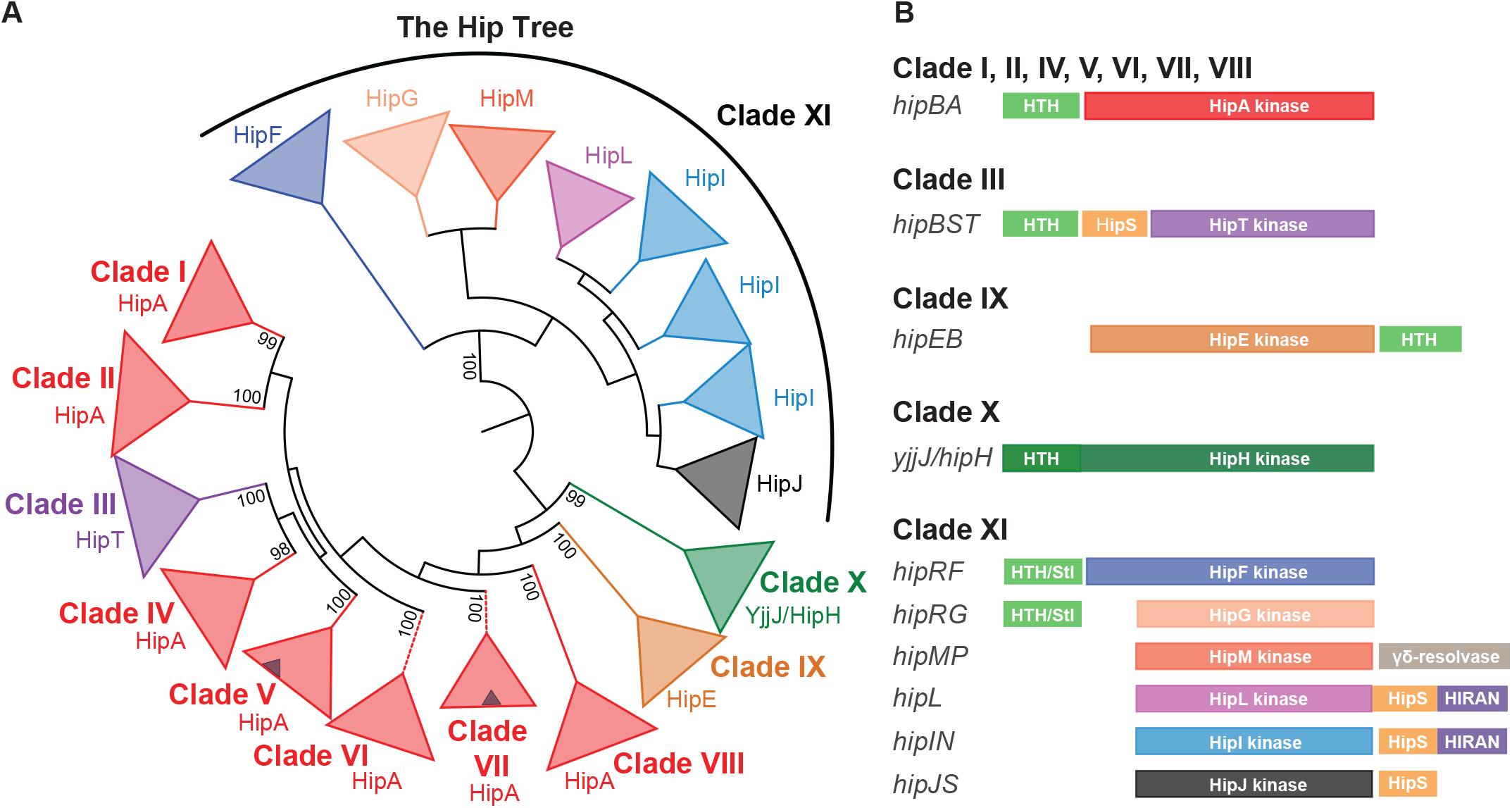
Phylogeny and genetic contexts of HipA-homologous kinases. (**A**) Simplified phylogenetic tree covering 1,239 HipA-homologous kinases (the “Hip Tree”, see **Figure S2** for details). The Hip Tree was divided into eleven Main Clades I to XI. The coloring of the Hip Tree reflects the genetic contexts that encoded the kinases such that each color corresponds to a distinct type of TA module. All main clades are monophyletic except Clade XI that consists of six different kinase families. Small blue triangles within the red triangles of Main Clades V and VII symbolize subclades of kinases encoded by TA modules with a reversed gene order relative to *hipBA* – that is – with the gene order *hipAB*. The Hip Tree was visualized by iTOL (82). (**B**) Genetic organizations of the TA modules encoding the 1,239 HipA-homologous kinases. The various types of genetic organization were obtained by manual inspection of the genes upstream and downstream of the kinase genes listed in **Table S1**. The coloring of the Hip kinases in (**B**) follows the coloring of the clades in (**A**). Putative antitoxins with Helix-Turn-Helix (HTH) domains are colored light green. Stl/HTH, putative antitoxins containing HTH domain and a domain with structural similarity to the “polyamorous” repressor Stl encoded by *Staphylococcus aureus*; HipS, HipS-like; HIRAN, HIP116 Rad5p N-terminal domain).

### HipA-homologous kinases are encoded by 10 different genetic organizations

By careful investigation of the sequence data set in **Table S1**, we found that a consistent and biological meaningful definition of “Hip kinase family” can be based on the genetic context encoding the kinases. Using this classification, we identified a total of 10 Hip kinase families encoded by TA modules with 10 different genetic organizations (**Figure 2B**). The frequencies of the TA modules with different genetic organizations are given in **Table S2B**.

Using this classification, all HipA kinases encoded by *hipBA* modules with an upstream HipB HTH antitoxin gene cluster together in Clades I, II, and IV to VIII. Experimentally characterized *hipBA* modules from *E. coli* K-12 and *C. crescentus* are all in Clade I (**Figure S2A**). Interestingly, Clades VI and VII each contain a subclade that have a reversed gene order (i.e. *hipAB*), indicating that gene reversion occurred independently several times during evolution (indicated as blue triangles within red triangles in **Figure 2A** and as branches with dashed lines in **Figure S2A**). Similarly, Clade IX consists entirely of kinases encoded by operons with a reversed gene order relative to *hipBA* and the deep branching of this clade warrants the definition of a new kinase family that we call HipE (**Figures 2A**). HipE kinases are encoded by *hipEB* operons that also encode putative HipB antitoxins with HTH domains (**Figure 2B**). Clade III consists solely of HipT kinases encoded by *hipBST* operons, including the characterized locus from *E. coli* O127, while Clade X consists entirely of YjjJ-like kinases encoded by monocistronic operons. Interestingly, Clade XI is highly diverse and contains no less than six kinase families (**Figure 2A**) encoded by different genetic contexts (**Figure 2B**). To standardize the nomenclature, all genes encoding putative antitoxins with HTH domains have been named HipB except for HipB of *hipBST* and HipR of *hipRF* and *hipRG*. In all cases investigated, HipB functions as both an antitoxin and an autoregulator of transcription with the exception of the HipB protein encoded by the *hipBST* operons, which autoregulates transcription but does not function directly as an antitoxin as mentioned above (20, 59). As described later HipR encoded *hipRF* and *hipRG* operons also contains a HTH domain but exhibits structural similarity with the Stl repressor.

### Conserved HipA kinases appear to contain functionally significant differences

Clade I contains the “classical” bacterial HipA kinases with known cellular targets, HipA of *E. coli* K-12, HipA_1_ and HipA_2_ of *C. crescentus*, together with their close homologs. An alignment of representative sequences of subclades containing HipA, HipA_1_ and HipA_2_ kinases reveals, as expected, the four canonical core kinase motifs: the Gly-rich loop, the Activation loop, the Catalytic motif, and the Mg^2+^-binding motif (**Figure S3**). The alignment also reveals a number of insertions (ω1 to ω6) and deletions (Δ1 to Δ3), also called “indels”, in the two HipA_*Ccr*_ subclades relative to the HipA_*E. coli* K-12_ subclade. **Figure 3A** shows a schematic overview of HipA indicating the positions of the indels relative to the core kinase motifs in the primary sequences, while **Figure 3B and 3C** show a mapping of the HipA_1_ and HipA_2_-specific indels as well as regions of high sequence divergence onto the structure of HipA of *E. coli* K-12. Interestingly, despite being distant in the primary sequence, the two largest deletions in *E. coli* HipA (Δ2 and Δ3) are adjacent in the tertiary structure (**Figure 3B and 3C**) and close to the γ-phosphate of ATP. Additionally, both *C. crescentus* HipA kinases share a C-terminal region of high sequence divergence that maps to solvent exposed residues of a surface helix (**Figure 3A**) while HipA_2_ has an additional unique region of divergence (**Figure 3C**). Studies of eukaryotic cyclin-dependent kinases have shown that the homologous region where Δ2, Δ3 and both regions of divergence are situated is involved in target binding (61), raising the possibility that the differences observed relate to differences in target specificity. Finally, we note that HipA_2_ has several regions (ω1”, ω2”, and ω3”) not present in HipA (**Figure 3A and S3**) concentrated in the region that interacts with the very C-terminus of HipB in the *E. coli* HipBA structure (**Figure 3C**) thus potentially affecting the mechanism by which the antitoxin interacts with and inhibits the cognate kinase.

**Figure 3.**
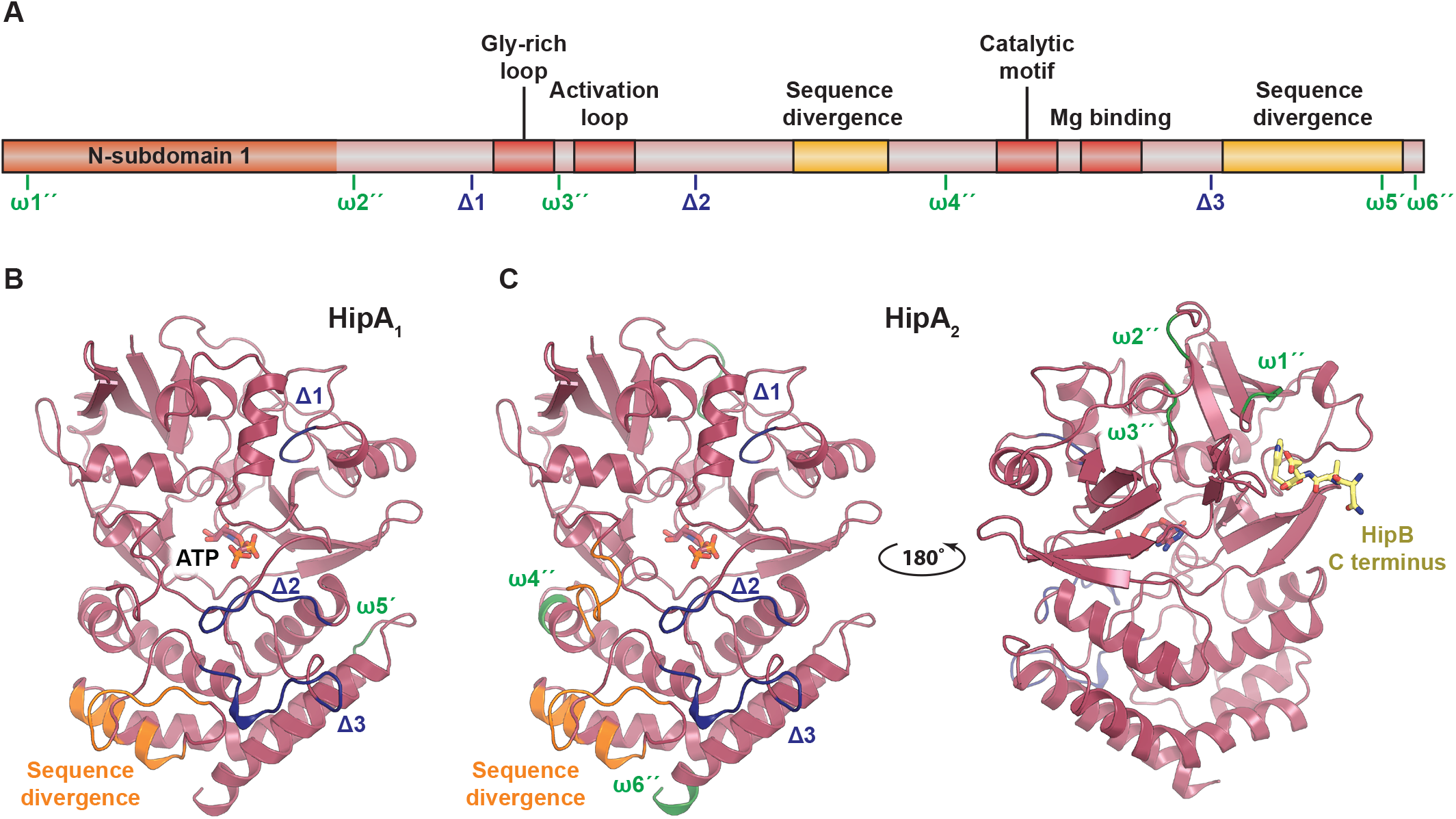
Structural mapping of insertions and deletions observed in characterized HipA kinases. (**A**) Schematic overview of HipA showing the conserved regions (the Gly-rich loop, Activation loop, Catalytic motif, and Mg binding motif) in red as well as insertions (ω, green) and deletions (Δ, blue) in *C. crescentus* HipA_1_ and HipA_2_ relative to HipA from *E. coli* K-12 based the sequence alignment shown in Figure S3. An insertion in HipA_1_ relative to HipA is denoted ω5’ while insertions in HipA_2_ relative to HipA are marked ω1’’, ω2’’, ω3’’, ω4’’, and ω6’’. Deletions in HipA_1_ and HipA_2_ relative to HipA are marked Δ1, Δ2, and Δ3. Regions of high sequence divergence are shown in yellow. (**B**) Mapping of the insertions and deletions in HipA_1_ onto the structure of HipA of *E. coli* K-12 (PDB: 3FBR) using the same nomenclature and color scheme as in (A) (45). ATP is shown with colored sticks. (**C**) Mapping of the insertions and deletions in HipA_2_ onto the structure of HipA. Note that the insertions ω1” and ω3” in HipA_2_ are located close to the region that in HipA interacts with a C-terminus of HipB (yellow sticks) and could perhaps affect antitoxin activity.

### All HipT kinases lack the canonical N-terminal subdomain

Main Clade III consists exclusively of HipT kinases (**Figure 2**). As shown before, the sequences of the HipT kinases of *E. coli* O127, *H. influenzae*, and *T. auensis* align co-linearly with the C-terminal part of *E. coli* K-12 HipA (**Figure S4A**), but lack the canonical ~100 aa N-subdomain-1 that serves as a lid on top of the core kinase domain in HipA (59). Nevertheless, the four conserved Hip kinase motifs (Gly-rich loop, Activation loop, Catalytic motif, and Mg^2+^-binding motif) are conserved in the HipTs, as well as a serine adjacent to the Gly-rich loop that is subject to auto-phosphorylation in both HipT and HipA (46, 59). We showed previously that HipS, which is encoded immediately upstream of HipT, functions as the antitoxin of *hipBST* modules and that this protein exhibits sequence similarity to the N-terminal domain of HipA (59) (**Figure S4A and S4B**). In other words, the “missing” N-subdomain-1 of HipT appears to be encoded immediately upstream and thus in a similar genomic location relative to the core kinase domain as in HipA, which suggests that the gene has been split (or merged) at some point during evolution. However, the functional and structural implications of this difference with respect to kinase activation and regulation, are not yet understood.

### YjjJ kinases contain an N-terminal HTH domain

YjjJ of *E. coli* K-12 is a HipA homolog encoded by a monocistronic operon and thus “lacking” a closely linked antitoxin or DNA-binding HipB-like gene. Instead, YjjJ has a ~100 aa N-terminal extension containing a Helix-Turn-Helix (HTH) domain (residues 15 to 34) that potentially could function as a *cis*-acting antitoxin, a property previously found with other Type II modules (62). To maintain a uniform nomenclature, we propose here to rename YjjJ kinases HipH (H for <u>H</u>TH domain) as these constitute the monophyletic Clade X in the Hip Tree (**Figure 2A**). To avoid alignment of non-homologous domains, we chose to align subclade HipH_*E. coli* K-12_ with HipT_*E. coli* O127_ because HipT lacks the ~100 aa N-terminal domain present in HipA but has all the canonical kinase domains. As seen from **Figure 4**, HipH and HipT kinases align co-linearly with respect to the four conserved kinase features while the N terminus of HipH forms a separate HTH domain. Moreover, HipH kinases contain two conserved serine residues near the Gly-rich loop, raising the possibility that they are regulated through auto-phosphorylation in a way similar to HipT (59).

**Figure 4.**
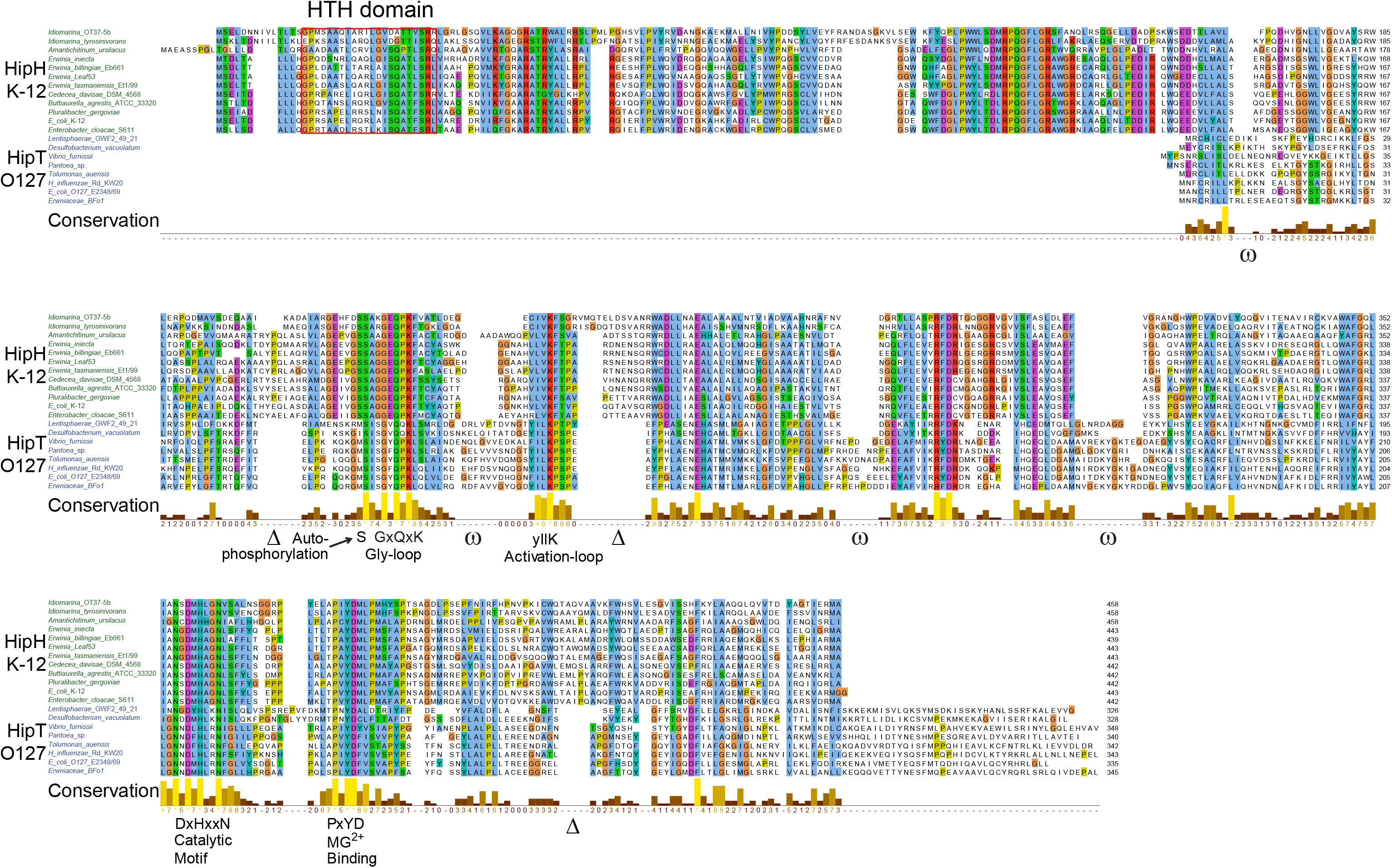
Alignment of HipH and HipT kinases. Sequence alignment of subclades containing HipT of *E. coli* O127 and HipH of *E. coli* K-12. Deletions (Δ) and insertions (ω) relative to the HipH_*E coli* K-12_ subclade are indicated, as well as the four conserved kinase motifs (Gly-loop, Activation-loop, Catalytic motif and Mg^2+^ binding motif). HTH domains in the N-terminus of HipH kinases are boxed in in red.

### Main Clade XI consists of kinases belonging to six different families

The kinases in Main Clade XI of the Hip Tree are encoded by six different genetic contexts and define six novel Hip kinase families (**Figure 2A, 2B**). All six subclades are separated by high bootstrap values, supporting that their separation into novel kinase families is phylogenetically justified (**Figure S5**). The six families encompass four families of short kinases (HipG, HipM, HipI, and HipJ), one longer variant (HipF) similar in size to HipA and a “long” kinase family (HipL) (**Figure 2B**). In the following, we describe these six new types of kinase-encoding TA modules.

### The HipG, HipM, HipI, and HipJ families contain the core kinase domain

The HipG, HipI, HipJ, and HipM kinases are all phylogenetically closely related and are, on average, even smaller than the HipT kinases (279-321 aa versus 291-346 aa, respectively) (**Table S1** and **Figure S5**). A sequence alignment of representatives of these families confirms that they contain the four conserved core Hip sequence motifs are thus likely active kinases (**Figure S6**). However, like HipT, these kinases lack the N-subdomain-1 present in HipA, HipE and HipF and consequently, the Gly-rich loop is located close to the N-termini (**Figure S6**). The kinases in the HipG group do not have a conserved serine or threonine adjacent to the Gly-rich loop, suggesting they are probably not regulated by auto-phosphorylation in a way similar to HipA. In contrast, members of the other families (HipM, HipI, and HipJ) mostly have either Ser or Thr near the Gly-rich loop, suggesting on the other hand that they may be regulated in this way. We note that HipI kinases constitute four separate subclades of the phylogenetic reconstruction of Main Clade XI (**Figure 2A**). Three of the subclades consists of kinases from Actinomycetes and Firmicutes while the fourth consists of a mix of kinases from proteobacteria and cyanobacteria (**Figure S5**).

### HipF and HipG kinases have putative antitoxins related to polyamorous repressor Stl

HipF and HipG kinases are encoded in operons with putative antitoxins that we call HipR (**Figure 2B**). Interestingly, Phyre2 (63) predicts HipR to be structurally related to the two-domain transcriptional repressor Stl encoded by *Staphylococcus aureus* superantigen-carrying pathogenicity islands (*Sa*PI). Stl has a HTH domain and maintains integration of *Sa*PI elements in the bacterial chromosome by transcriptionally repressing genes essential for element excision and replication (64). *Sa*PI excision and replication is induced by invading phages via specific interaction between Stl and non-essential phage proteins (65–69). These observations allow us to hypothesize that phage proteins could potentially activate HipF and/or HipG kinases during infection via interaction with HipR. Under this model, activation of the kinases induces abortive phage infection by inhibition of translation via phosphorylation of an essential component of the protein synthesis apparatus, thus eliciting phage resistance. This possibility can now be tested experimentally.

### Putative HipN antitoxins consist of a HipS-like and a HIRAN domain

The genes encoding HipM, HipI and HipJ kinases all have downstream genes coding for putative antitoxins (**Figure 2B**). The putative antitoxin HipN encoded by *hipIN* is a two-domain protein consisting of an N-terminal HipS-like (**Figure S7A**) and a C-terminal HIRAN domain (**Figure S8A**). The HipS-like domain of HipN may function to neutralize its cognate HipI kinase as is the case of HipS encoded by *hipBST* (59). A possible function of the HIRAN domain of HipN is discussed below.

### A second family of putative HipS antitoxins

*hipJS* modules encode a HipJ kinase and a putative antitoxin HipS that exhibits similarity to HipS of *hipBST* (**Figure S7A**). Thus, similarly to HipT (59), HipJ may be neutralized by its cognate HipS. As noted above, HipS and HipS-like domains exhibit sequence similarity with the ~100 aa N-subdomain-1 of HipA and this domain was therefore included in the alignment of the HipS antitoxins and HipS-like domains (**Figure S7A**). The phylogenetic tree based on the HipS and HipS-like sequences is bifurcated, with HipS encoded by *hipBST* and *hipJS* branch together with N-subdomain-1 of HipA while the HipS-like sequences encoded by *hipL* and *hipN* generate a distinct second branch (**Figure S7B**). HipJ kinases are closely related to HipI kinases (**Figure S5**) and it is tempting to speculate that *hipJS* modules evolved from *hipIN* modules by deletion of the HIRAN domain of HipN antitoxins (**Figure 2B**).

### HipL kinases contain both HipS-like and HIRAN domains

HipL kinases, encoded by monocistronic operons, form a single subclade of Main Clade XI that is further divided into two subclades (**Figure S5**). The two subclades consist of HipL kinases from Gram-positive and Gram-negative bacteria, respectively. Their sequences are clearly distinct, with many subclade-specific insertions and deletions in the kinase core domain (**Figure S8B**). Alignment of HipL and HipA reveals that HipL maintains the four, conserved core kinase motifs and has a large C-terminal extension of ~200 aa of which the ~100 aa at the extreme C-terminus are annotated at GenBank as a HIRAN domain (**Figure S8C**). Like the small kinases, HipL kinases lack the N-subdomain-1 of ~100 aa relative to the core kinase domain of HipA (**Figure S8C**). Unexpectedly, however, the ~100 aa domain located between the core kinase domain and the C-terminal HIRAN-domain exhibits sequence similarity with HipS (**Figure S8D**). The N-terminal sequences of HipA also align well with the HipS-like domain of HipL (**Figure S8E**), consistent with the fact that HipS exhibits sequence similarity with the N-subdomain-1 of HipA (**Figure S4B**). In summary, HipL kinases are three-domain proteins consisting of an N-terminal kinase core domain, a HipS-like domain and a C-terminal HIRAN domain (**Figures 2B** and **S9B**). We note the possibility that the HipS-like domain may be involved in regulating the kinase activity of HipL, reminiscent of how HipS regulates the kinase activity of HipT (59).

### The HIRAN domains of HipL and HipN may bind DNA

The presence of HIRAN domains in both HipL kinase and the putative antitoxins HipN is interesting, not least because HIRAN domains can bind DNA. Moreover, the HIRAN domains of HipL kinases and HipN antitoxins are in both cases joined with a HipS-like domain (**Figure 2B**). HIRAN domains have previously been identified in eukaryotic multi-domain DNA repair proteins (70). Experimentally analyzed HIRAN domains bind single-stranded or double-stranded DNA ends (71, 72). Structural studies of human HLTF, a DNA helicase implicated in remodelling of replication forks, including fork regression and restart (73), revealed the residues required for DNA binding (72) (**Figure 5A, 5B**). Due to a high sequence divergence, we decided to split the HIRAN domains of the HipN homologues based on their phylogenetic origin (**Figures 5C** and **S8**). Most of the HIRAN domains retained the majority of the sequence motifs that interact with DNA, with the exception of HipN from Actinomycetes. A study of the HIRAN-domain of human HLTF showed that Phe142 (NA<u>F</u>) is required for binding to duplex DNA because it stacks with nucleobases of the other strand (72) (**Figure 5B**). Importantly, most of the bacterial HIRAN domains lack a conserved Phe at this position (**Figure 5C**), raising the possibility that the HIRAN domains in HipN and HipL interact with ssDNA.

**Figure 5.**
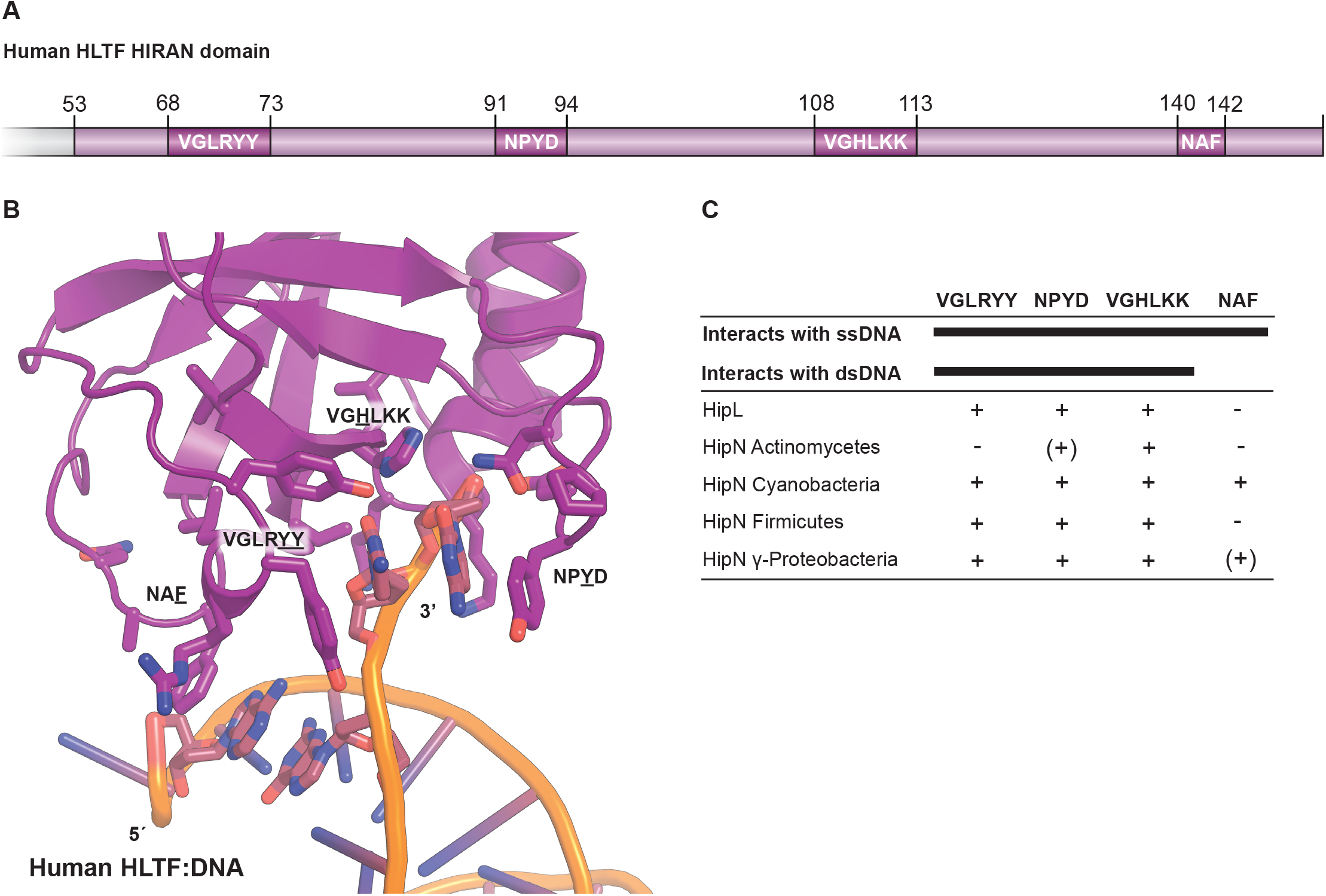
Comparative analysis of HIRAN domains. The HIRAN domains of HipL and HipN have sequence motifs required for DNA-binding. (**A**) Schematic overview of the HIRAN domain found in human Helicase-like transcription factor (HLTF) showing the relative positions and sequences involved in ssDNA and dsDNA binding (72). (**B**) Structure of the human HLTF HIRAN domain with the four regions necessary for DNA binding shown as sticks (PDB: 4XZF) (72). (**C**) Overview of sequence motifs present (+) or absent (−) in homologues of HipL and HipN.

The function(s) of the HIRAN domains of HipL and HipN are, of course, unknown. One attractive possibility is that the HIRAN domains of HipL and HipN recognize DNA ends (ssDNA or dsDNA, or both) of invading phages or phage DNA replicative intermediates that, in turn, leads to activation of the HipL and HipI kinases. Activation of the kinases could lead to phosphorylation and inhibition of an essential cellular component, thereby triggering abortive phage infection. Abortive phage infection is a common biological function of TA modules (74, 75) and the possibility that *hipL* and *hipIN* function to curb phage infection can now be tested experimentally.

### A putative antitoxin with a γδ resolvase-like domain

A final variation among the Hip kinases is found for the *hipMP* modules, which encode a HipM kinase that is short and closely related to HipG kinases (**Figure 2A**) and a putative HipP antitoxin that exhibits similarity to γδ-resolvases (**Figure 2B**). Bacterial γδ-resolvases are transposon-encoded enzymes that catalyze recombination within a complex nucleoprotein structure during site-specific DNA recombination and thus have the capability to bind DNA (76). Interestingly, γδ-resolvases are structurally related to 5’-3’ exonucleases active on RNA, which may also provide clues to the role of this domain in the context of Hip kinases (77). By analogy with other *hipBA* modules (28, 78), we therefore postulate that HipP functions as antitoxin in the *hipMP* systems and may bind DNA to autoregulate transcription as seen for many type II TA systems.

## Conclusion

In addition to the three known kinase families HipA, HipT and YjjJ/HipH, we discover here seven novel Hip kinase families encoded by different genetic contexts. Kinases of one family, HipE, are encoded by TA operons with a reversed gene order relative to *hipBA*, while kinases of families HipF and HipG have putative Stl-homologous antitoxins that may be regulated by proteins encoded by attacking phages. Kinases of one family, HipJ, are associated with HipS-domain putative antitoxins that, similar to HipS of *hipBST*, may interact with and neutralize their cognate kinase toxins. Kinases of yet another family, HipI, are associated with putative antitoxins that consist of a HipS-like domain and a HIRAN DNA-binding domain while HipJ kinases are associated with putative antitoxins exhibiting similarity with γδ-resolvases. Finally, HipL kinases, encoded by monocistronic operons, consist of an N-terminal core kinase domain, an internal HipS-like domain and a C-terminal HIRAN DNA-binding domain. The latter two domains of HipL may function in regulating its kinase activity. Our analysis presented here builds a foundation for the future experimental analysis of HipA-homologous kinases.

## Acknowledgements

We thank Boris Macek (Proteome Center Tübingen, University of Tübingen, Germany) and Yong E. Zhang (Department of Biology, University of Copenhagen, Denmark) for critical comments to the manuscript.

## Methods

### Data sampling

The sequences of Hip toxins with known cellular targets as of December 2019 were HipA and HipH/YjjJ of *E. coli* K-12 and HipT of *E. coli* O127. These sequences were used as seeds in BLASTP searches at https://blast.ncbi.nlm.nih.gov/, using the bacterial phyla shown in **Figure S1** as search spaces. HMMSEARCH at https://ebi.ac.uk (79) was used to expand poorly populated Clades. In total, ≈1,800 Hip sequences were retrieved (E > 10^−10^) and curated manually such that every Hip kinase sequence retained satisfied the following criteria: (i) the Hip kinase gene should encode a full-length kinase with the four canonical kinase motifs (Gly-rich loop, Activation loop, Catalytic motif and Mg^2+^ binding domain), as determined from a multiple sequence alignment (MSA); (ii) kept kinases should be encoded by a gene with an adjacent upstream or downstream putative antitoxin gene; (iii) in general, the adjacent, putative antitoxin gene should encode a DNA-binding protein (although this criterion is not satisfied by HipF proteins); In **Table S1**, kept kinases are less than 95% identical to any other kept kinase. By this scrutiny, the initial gene set was reduced to 1,239 Hip gene modules (**Table S1** and **Figure 2**)

### Toxin – antitoxin module gene organization

The gene organizations shown in **Figure 1** were deduced by manual inspection of genes neighboring the Hip kinase genes of **Table S1**.

### Sequence alignments and phylogenetic tree reconstruction

Sequence alignments were was generated by Clustal Omega (80) at https://www.ebi.ac.uk and imported into Jalview (81). Protein sequence alignments in Jalview 2.11.0 were exported as vector files (EPS or SVG formats) and imported into Adobe Illustrator CS6, annotated and saved in PDF format for publication. Phylogenetic trees were visualized using iTOL (82). Reconstruction of phylogenetic trees was accomplished using IQ-TREE that uses the Maximum Likelihood approach and Ultrafast bootstrapping via the CIPRES module in Genious Prime (83–85). The kinase sequence alignments used in the reconstruction of the 3 phylogenetic trees that we present had the following characteristics: (1) The sequence alignment of the 1,239 kinases (**Figure S2**) has 1556 columns, 1511 distinct patterns, 1172 parsimony-informative, 172 singleton sites and 211 constant sites; (2) The sequence alignment of the 81 sequences of Main Clade XI (**Figure S5**) has 785 columns, 761 distinct patterns, 650 parsimony-informative, 58 singleton sites and 77 constant sites; (3) The sequence alignment of the 112 HipS and HipS-like sequences (**Figure S7B**) has 170 columns, 170 distinct patterns, 146 parsimony-informative, 16 singleton sites and 8 constant sites. The alignment of the HipS and HipS-like sequences has fewer singleton and constant sites thus explaining the low bootstrap values (**Figure S7B**).

Structure similarity searches was done using Phyre2 (63) (http://www.sbg.bio.ic.ac.uk/phyre2/) and mapping of deletions and insertions on existing structures using PyMOL. HTH motifs were identified by two different algorithms, EMBOSS (86) and HELIX-TURN-HELIX MOTIF PREDICTION (87).

## Legends to Supplemental Figures and Tables.

**Figure S1. Hip kinases divided on the Tree-of-Life**. Overview of prokaryotic phyla used as search spaces in BLASTP and HMMSEARCH procedures. HipA and HipH/YjjJ of *E. coli* K-12 and HipT of *E. coli* O127 were used as queries in BLASTP and HMMSEARCH searches using individual prokaryotic phyla as search spaces. The colored dots reveal the frequencies of Hip homologues of the individual phyla included in the analysis and are indicated in the insert. Kinase frequencies of individual phyla were collapsed to maintain clarity while the exact numbers are given in **Table S2A**. The figure was adapted from Brock Biology of Microorganisms (15^th^ Ed).

**Figure S2. Phylogenetic Tree of 1,239 HipA-homologous kinases**. (**A**) The Hip Tree of Hip kinases annotated with phylogenetic origin. Roman numerals I to XI denote the eleven deep-branching Main Clades coloured as in **Figure 2**. Kinase labels reveal phylum, gene size (in codons) and GenBank identifier (from **Table S1**). Stippled branches in Main Clades V and VII indicate TA operons with a reversed gene order relative to *hipBA*. The positions of HipA of *E. coli*, HipA_1_, HipA_2_ and HipA_3_ of *C. crescentus*, HipA of *S. oneidensis* MR-1, HipT of *E. coli* O127 and YjjJ (HipH) of E. coli K-12 that have been investigated experimentally and are indicated. The outer ring shows the phylogenetic origin of the individual kinases in color with the codes: proteobacteria: blue; Actinomycetes: green; Firmicutes: red; Planctomycetes, Verrucomicrobia, and Chlamydiae (PVC) group: orange; Archaea (Crenarchaeota; Euryarchaeota and candidatus Woesearchaeota); Spirochaetes: light green; Bacteroidetes: Magenta; Cyanobacteria: light cyan; Deferribacteraceae: brown; Acicdobacteria: black; Fibrobacter: light grey; Aquificae and Thermotogae: dark grey; all other phyla (Nitrospirae, Tenericutes, Fusobacteria, Chlorobi, Ignavibacteriae and Chrysiogenales): dark green. Description of the Main Clades: **Clade I** consists of 712 kinases all encoded by *hipBA* modules and encompassing HipA of *E. coli* K-12, HipA_1_ and HipA_2_ of *C. crescentus*, and HipA of *S. oneidensis* that have been investigated experimentally (**Table 1**). **Clade II** consists of 73 kinases, also encoded by *hipBA* loci but is distinct from Clade I by its deep branching and the high bootstrap value (100%) that separates the two clades. Clade II is dominated by kinases from Actinobacteria. **Clade III** consists of all 48 HipT kinases encoded by the tricistronic *hipBST* modules. **Clade IV** consists of 14 kinases, all encoded by *hipBA* modules. Clade IV kinases are all from Proteobacteria. **Clade V** consists of 9 kinases, all encoded by *hipBA* modules. Most of these kinases are from Proteobacteria. **Clade VI** consists of 36 kinases, 31 of which are encoded by canonical *hipBA* operons while 5 are encoded by operons with a reversed gene order (i.e. *hipBA*). **Clade VII** consists of 136 kinases encoded by 133 *hipBA* operons while 3 of the kinases are encoded by operons with a reversed gene order (i.e. *hipAB*). Clade VII kinases are encoded by a heterogenous set of organisms encompassing Proteobacteria, the PVC group, Acidobacteria, Spirochaetes, Tenericutes and Archaea. **Clade VIII** consists of 21 kinases, all encoded by *hipBA* modules. Most of Clade VIII kinases are from Proteobacteria while a few are from Nitrospirae and Archaea. **Clade IX** consists of 12 HipE kinases all encoded by operons *hipEB* operons with a reversed gene order relative to *hipBA* (**Figure 2B**). **Clade X** consists of 101 YjjJ/HipH kinases, all encoded by monocistronic operons. The majority of the YjjJ/HipH kinases are from Proteobacteria while a few are from the PVC group or Chrysiogenales. Clade XI, the most complex clade, consists of 81 kinases encoded by six different genetic contexts and contains six novel kinase families (**Figure 2B**). (**B**) Bootstrapped version of the Hip Tree shown in (A). The Hip Tree is based on a sequence alignment of the 1,239 kinases listed in **Table S1** and was generated by Clustal Omega (89), bootstrapped by IQ-TREE (84) via a module provided by CIPRES (http://www.phylo.org/) in Genious Prime and visualized by iTOL (82). The best-fit model chosen for phylogenetic tree reconstruction was WAG+F+R10 according to the Baysian Information Criterion (BIC).

**Table 1.** Overview of the Clades of the Hip Tree consisting of 1,239 Kinases. Data compiled from **Table S1**.

| Clade | Number of kinase genes | TA Gene Organization<sup>a)</sup> | Experimentally Analyzed TA Modules | Cellular Target(s) of Toxin | GenBank ID<sup>b)</sup> | References |
| --- | --- | --- | --- | --- | --- | --- |
| **I** | 712 | *hipBA* | *hipBA* of *E. coli* K-12<br>*hipBA*<sub>1</sub> of *C. crescentus*<br>*hipBA*<sub>2</sub> of *C. crescentus*<br>*hipBA*<sub>3</sub> of *C. crescentus*<br>*hipBA* of *S. oneidensis* MR-1 | GltX<br>Unknown or GltX and LysS<sup>d)</sup><br>LysS or GltX, LysS and TrpS<sup>d)</sup><br>Unknown<sup>d)</sup><br>Unknown | NP_416024.1(440aa)<br>ACL93947.1(423aa)<br>ACL96286.1(444aa)<br>WP_010920611.1(435aa)<sup>c)</sup><br>AAN53784.1(433aa) | (47, 51, 52, 57, 58) |
| **II** | 73 | *hipBA* | None |  |  |  |
| **III** | 48 | *hipBA* | *hipBST* of *E. coli* O127<br>*H. influenzae* KW20<br>*T. auensis* DSM9187 | TrpS<br>TrpS<br>TrpS | WP_015879003.1(342aa)<br>NP_438824.1(343aa)<br>CAS11333.1(335aa) | (59) |
| **IV** | 15 | *hipBA* | None |  |  |  |
| **V** | 19 | *hipBA* | None |  |  |  |
| **VI** | 36 | *hipBA* & *hipAB* | None |  |  |  |
| **VII** | 132 | *hipAB* & *hipAB* | None |  |  |  |
| **VIII** | 21 | *hipBA* | None |  |  |  |
| **IX** | 12 | *hipEB* | None |  |  |  |
| **X** | 101 | *hipH/yjjJ* | *hipH/yjjJ* |  | P39410.1 (443aa) | (60) |
| **XI** | 81 | *hipRF*, *hipRG*,<br>*hipMP*, *hipL*,<br>*hipIN* & *hipJS* | None |  |  |  |

a) Gene organization refers to information retrieved from **Table S1** and visualized schematically in **Figure 2B**
b) GenBank IDs of the kinases are from **Table S1** and their position in the Hip Tree visualized in **Figure S2A and S2B**
c) *hipBA*<sub>3</sub> of *C. crescentus* previously had the accession number NP\_421566.1 and was used in **Table S1**
d) The substrate of HipA<sub>1</sub> is either unknown (58) or GltX plus LysS (57) while the substrate of HipA<sub>2</sub> is either TrpS (58) or GltX, LysS plus TrpS (57). The substrate of HipA<sub>3</sub> is unknown.

**Figure S3. Alignment of HipA kinases that have different cellular targets**. Sequences of subclades containing HipA of *E. coli* K-12, HipA_1_ and HipA_2_ of *C. crescentus* NA1000 were aligned. The subclades were selected such that they are separated by high bootstrap values (**Figure S2**). Deletions (Δ) and insertions (ω) relative to the HipA_*E. coli* K-12_ subclade are indicated as well as the four conserved kinase motifs (Gly-loop, Activation-loop, Catalytic motif and Mg^2+^ binding motif). Clade-specific insertions in HipA_1_ and HipA_2_ are indicated with red boxes.

**Figure S4. HipT and HipS exhibit sequence similarity with the C and N-terminal ends of HipA, respectively**. (**A**) Subclade HipA_*E. coli* K-12_ of Main Clade I aligned with subclade HipT_*E. coli* O127_ of Main Clade III (**Figure S2**). The four conserved canonical Hip kinase motifs are indicated as well as a conserved, autophosphorylated serine residue. Conserved deletions (Δ) and insertions (ω) in the HipT kinases relative to the HipA kinases are also indicated. (**B**) Subclade HipA_*E. coli* K-12_ aligned with subclade HipS_*E. coli* O127_ antitoxins.

**Figure S5. Phylogenetic Tree of Main Clade XI**. The clade consists of 81 kinases belonging to the six novel Hip families HipF, HipG, HipI, HipJ, HipM and HipL. HipF and HipL kinases (encoded by *hipRF* and *hipL* loci) are located to single subclades while HipG, HipI, HipJ and HipM are located to four different subclades where the kinases are encoded by TA modules with different genetic organizations (*hipRG*, *hipIN*, *hipJS* and *hipMP*). The genetic organizations are shown schematically in **Figure 2B**. The HipG, HipI, HipJ and HipM kinases have similar sizes and are phylogenetically related; however, the bootstrap values and the subclade-specific genetic organizations warrant their designation as separate kinase families. The best-fit model chosen for phylogenetic tree reconstruction was LG+F+R5 according to BIC.

**Figure S6. Alignment of the HipG, HipI, HipJ and HipM kinases of Clade XI**. The four conserved Hip kinase motifs are indicated. The kinases of these 4 families are encoded by TA modules with 4 different genetic organizations (**Figure 2B**) and have very short N-terminal domains.

**Figure S7. Sequence alignments and phylogeny of HipS and HipS-like domains**. (**A**) Alignment of HipS encoded by *hipBST* modules with HipS-like domains of HipN encoded by *hipIN*, HipS encoded by *hipJS* and HipS-like domain encoded by *hipL*. For comparison, the N-terminus of HipA was included in the alignment. (**B**) Phylogenetic tree derived from the sequence alignment shown in (A). **The best-fit model chosen for reconstruction of the phylogenetic tree was LG+R5 according to BIC**.

**Figure S8. Alignment of HIRAN domains and of HipL Kinases**. (**A**) The HIRAN domain of human HLTF aligned with the HIRAN domains of HipL kinases and putative HipN antitoxins. The sequence motifs of the HIRAN domain of HLTF involved in interaction with DNA are shown in boxes and are color-coded (see also **Figure 5**). The HipN sequences were divided into 4 separate alignments according to phylogeny in order to reveal the similarities with the sequence motifs of the HIRAN domain of HLTF. Color-codes of the individual sequence motifs: Black indicates conserved motifs that may have the capability to interact with DNA; red indicates that conserved motifs were not identified; orange indicates some conservation. (**B**) Alignment of the HipL kinases of Main Clade XI. The four core kinase motifs are indicated. The alignment reveals two subclades consisting of kinases from Gram-positive and Gram-negative species, respectively (also apparent from Figure S5). (**C**) Alignment of HipL kinases with subclade HipA_*E. coli* K-12_. Deletions (Δ) and insertions (ω) in HipL relative to HipA, as well as the core kinase motifs are indicated. (**D**) Alignment of HipL kinases with HipS antitoxin sequences (subclade HipS_*E. coli* O127_). As seen, the internal domain of HipL exhibits sequence similarity with HipS. (**E**) Alignment of HipL kinases with the N-terminal domains of HipA_*E. coli* K-12_ kinase subclade.

****Table S1**.Primary information on 1,239 HipA-homologous kinases**. The columns A to P yield the following information: (**A**) GenBank ID of kinase toxins; (**B**) Kinase gene lengths in codons; (**C**) Organism; (**D**) Label yielding phylum, gene length in codons and GenBank ID used the Hip Tree (**Figure S2**); (**E**) Kinase sequences; (**F**) Distance between kinase toxin and its cognate antitoxin in nucleotides (a minus indicates gene overlap); (**G**) Genetic organization (GO) of each toxin – antitoxin module; (**H**) Antitoxin GenBank ID; (**I**) Antitoxin gene length in codons; (**J**) Label yielding antitoxin phylum, gene length and GenBank ID; (**K**) Antitoxin sequence including HipBs encoded by *hipBST* loci (HipS antitoxin sequences have their own dedicated columns in the Table); (**L**)Label yielding HipS phylum, gene length and GenBank ID (used in HipS alignments of **Figure S4B, Figure S7A, Figure S8D**); (**M**) Distance between *hipS* and *hipT* in nucleotides; (**N**) HipS antitoxin sequences; (**O**) Domains in antitoxins; (**P**) Comments to individual entries in the Table. GO, Genetic Organization. Full names of phyla are given in **Table S2A**.

****Table S2**A. The 1,239 Hip kinases Divided on Phyla**. The numbers were extracted from the data in **Table S1**.

****Table S2**B. Frequencies of gene organizations of the 1,239 Hip Tree**. Gene organizations of TA modules encoding Hip kinase toxins and their frequencies summarized from **Table S1**.

